# Role of Strong Localized vs. Weak Distributed Interactions in Disordered Protein Phase Separation

**DOI:** 10.1101/2023.01.27.525976

**Authors:** Shiv Rekhi, Dinesh Sundaravadivelu Devarajan, Michael P. Howard, Young C. Kim, Arash Nikoubashman, Jeetain Mittal

## Abstract

Interaction strength and localization are critical parameters controlling the single-chain and condensed-state properties of intrinsically disordered proteins (IDPs). Here, we decipher these relationships using coarse-grained heteropolymers comprised of hydrophobic (H) and polar (P) monomers as model IDPs. We systematically vary the fraction of P monomers *X*_P_ and employ two distinct particle-based models that include either strong localized attractions between only H–H pairs (HP model) or weak distributed attractions between both H–H and H–P pairs (HP+ model). To compare different sequences and models, we first carefully tune the attraction strength for all sequences to match the single-chain radius of gyration. Interestingly, we find that this procedure produces similar conformational ensembles, nonbonded potential energies, and chain-level dynamics for single chains of almost all sequences in both models, with some deviations for the HP model at large *X*_P_. However, we observe a surprisingly rich phase behavior for the sequences in both models that deviates from the expectation that similarity at the single-chain level will translate to similar phase-separation propensity. Coexistence between dilute and dense phases is only observed up to a model-dependent *X*_P_ despite the presence of favorable interchain interactions, which we quantify using the second virial coefficient. Instead, the limited number of attractive sites (H monomers) leads to the self-assembly of finite-sized clusters of different sizes depending on *X*_P_. Our findings strongly suggest that models with distributed interactions favor the formation of liquid-like condensates over a much larger range of sequence compositions compared to models with localized interactions.

## Introduction

Concentrated assemblies of biomolecules that exist without a lipid membrane, termed membraneless organelles (MLOs) or biomolecular condensates, play a key role in several cellular processes that include cell organization,^1^ signaling, ^2^ and stress response^3^ in addition to their relevance in pathological studies.^4^ In general, MLOs have liquid-like characteristics and form *via* phase separation. ^1,5,6^ It has been found that intrinsically disordered proteins (IDPs) or intrinsically disordered regions (IDRs) within proteins primarily drive phase separation.^7,8^ A common hypothesis for the prevalence of IDPs and IDRs in MLOs is their ability to form multivalent contacts^7,9^ *via* hydrogen bonding,^10^ electrostatic interactions, ^11,12^ cation−*π* interactions^13,14^ as well as sp^2^/*π* interactions.^15,16^ However, quantifying the relative contributions of these interaction types to the phase separation process is non-trivial, thus impeding the accurate prediction of the phase behavior of proteins based on sequence.^9,17^

Despite these challenges, significant progress has been made in elucidating the sequence determinants of phase separation through experimental and computational studies. ^10–22^ Among these studies, the work on the low-complexity domain (LCD) of hnRNPA1^21,23^ and the FUS prion-like domain^22^ have shown that the critical temperature *T*_c_ and the saturation (dilute-phase) concentration *c*_sat_ were altered when the number of aromatic residues (tyrosine) was varied. In addition, the mutation of arginine to lysine while maintaining the tyrosine content in FUS-like proteins led to a significant increase in *c*_sat_, thus implying a reduction in phase-separation propensity.^13,14^ Further work on the FUS-LCD, using a combination of nuclear Overhauser experiments and all-atom molecular simulations, demonstrated that residues such as serine, glycine, and glutamine also play an important role in the stabilization of liquid-like condensates through interactions with other residues in the sequence. ^18,24^ Studies involving shuffling of charges in LAF-1 RGG^11^ and DDX4^12^ have shown that charge patterning, frequently quantified by the sequence charge decoration parameter,^25^ influences the phase-separation propensity of a protein, highlighting the key role played by electrostatic interactions. These findings have led to different interpretations of the interaction scenarios governing the phase separation of proteins.

One class of models considers only a few residue types responsible for phase separation and introduces strong localized interactions between certain amino acid pairs such as tyrosine–tyrosine or tyrosine–arginine.^7,13,14,26^ Such models have been used to investigate single-component and multi-component phase separation. These models are in the spirit of the HP model popularized by Dill and coworkers to investigate protein folding, in which the protein is composed of hydrophobic (H) and polar (P) monomers, and there are energetically favorable interactions (attractions) between only the H monomers.^27–29^ More recent simulations using an off-lattice HP variant studied the different assemblies that formed when varying the fraction of H monomers at fixed chain length and fixed interaction strength.^30^

Another class of protein models considers the collective contribution of all amino acids in a protein sequence by employing weak distributed interactions, comprised of various interaction modes, distributed along the protein.^10,18,31^ Transferable coarse-grained models developed by some of us, using amino acid specific contact potentials based on hydrophobicity, are reflective of this consideration and have been used to successfully characterize the phase behavior of reference proteins like FUS-LCD, LAF-1 RGG, *α*-synuclein, and A1-LCD.^19,23,32–34^ An extension of the model applying an additional temperature-dependent parabolic scaling for the nonbonded interactions was shown to capture experimentally observed ^35^ upper critical or lower critical solution temperature transitions for a set of thermoresponsive model IDPs. ^36^

In this work, we investigate the differences between these two classes of models by systematically studying the phase behavior of a coarse-grained heteropolymer consisting of H and P monomers. We consider an off-lattice variant of the HP model, which has strong localized attractions between only H monomers, and an extension we call the HP+ model, which has weak distributed attractions between both H–H and H–P monomer pairs. To make the different sequences and models comparable, we carefully scale the interactions between H monomers for each sequence to achieve the same single-chain dimensions as a purely hydrophobic reference sequence. Despite similarity at the single-chain level, differences between sequences and models emerge in the condensed-state, emphasizing the importance of the nature of intra- and interchain interactions in the formation of a homogeneous liquid-like condensate. The implications of our findings are of particular interest to the experimental interpretation of distinct interaction scenarios driving the phase separation of a given disordered protein.

## Molecular Model and Simulation Methodology

We used a coarse-grained polymer model with a single bead per residue (monomer) and the solvent modeled implicitly in the effective monomer–monomer pair interactions. The sequences comprised of two types of monomers, namely hydrophobic (H) and polar (P) monomers having the same diameter *σ* = 0.5 nm and mass 100 g/mol. We fixed the chain length to *N* = 20 and randomly generated 21 sequences with the fraction of polar beads *X*_P_ varying from purely hydrophobic, *X*_P_ = 0, to purely hydrophilic, *X*_P_ = 1 (Fig. 1a). In addition to our primary sequence set, we also tested three other randomly generated sequence sets (RS1, RS2, and RS3 in Fig. S1) as well as a highly patterned sequence set (RS4 in Fig. S1).

**Figure 1:**
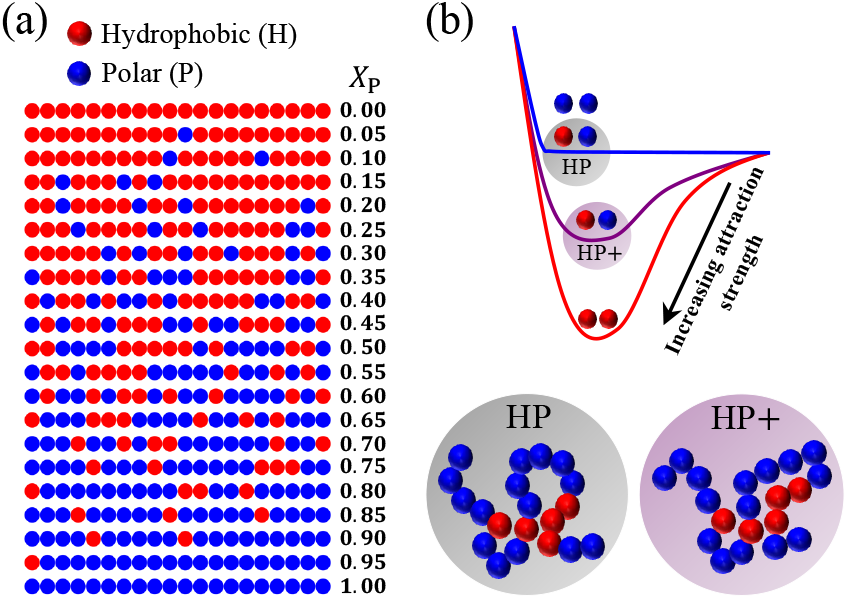
(a) Simulated sequences of length *N* = 20 with the fraction of polar beads *X*_P_ ranging from *X*_P_ = 0 (purely hydrophobic) to *X*_P_ = 1 (purely hydrophilic). (b) Schematic highlighting the different interactions used in the HP and HP+ models.

Interactions between bonded beads were modeled using a harmonic potential,

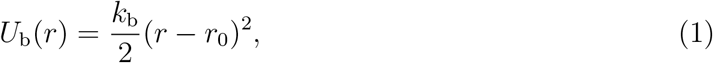

with distance *r* between monomers, spring constant *k*_b_ = 2000 kcal/(mol nm^2^), and equilibrium bond length *r*_0_ = 0.38 nm. Nonbonded interactions were modeled using a modified Lennard-Jones potential where the attractive contribution was scaled independently of the short-range repulsion by the pairwise hydropathy *λ_ij_* for monomers of type *i* and *j*,^19,37–39^

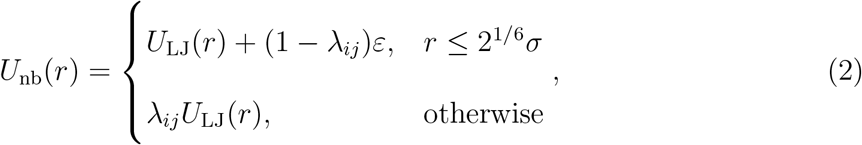

where *U*_LJ_ is the standard Lennard-Jones potential,

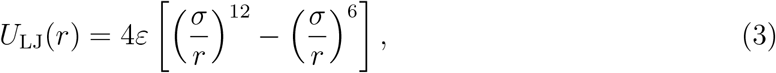

and *ε* is the interaction strength. We note that Eq. (2) reduces to the purely repulsive Weeks–Chandler–Andersen potential, ^39^ which describes excluded volume interactions between monomers under good solvent conditions, for *λ_ij_* = 0 and to Eq. (3), which also includes effective attraction between monomers as in a lower quality solvent, for *λ_ij_* = 1.

To determine *λ_ij_*, we accordingly assigned a nominal hydropathy *λ*_P_ = 0 to the P monomers and a nonzero hydropathy *λ*_H_ to the H monomers. (The specific value of *λ*_H_ was varied as described later.) We set *λ*_PP_ = *λ*_P_ and *λ*_HH_ = *λ*_H_ so that H–H monomer pairs were attracted to each other but P–P monomer pairs were not. We then proposed two different models (HP and HP+) for the cross-interactions between H–P monomer pairs controlled by *λ*_HP_. The HP model has strong localized attractions between only a certain type of residue, *e.g*., self-interactions between aromatic residues like tyrosine, so we set *λ*_HP_ = 0. The HP+ model also has weak distributed attractions with additional residues, *e.g*., cross interactions between aromatic residues and polar residues like glutamine, asparagine, serine, etc. (Fig. 1b), so we set *λ*_HP_ = (*λ*_H_ + *λ*_P_)/2 = *λ*_H_/2 to the mean hydropathy of the H and P monomers. In both models, we set *ε* = 0.2 kcal/mol, which, in conjunction with other model parameters, was previously shown to accurately capture IDP properties like radius of gyration *R*_g_.^19,33^

We performed Langevin dynamics simulations in the low-friction limit [friction coefficient 0.1g/(mol fs)] at fixed temperature *T* = 300 K for a total duration of 1 *μ*s using a timestep of 10 fs. We performed single-chain simulations using LAMMPS (29 October 2020 version), ^40^ while condensed-phase simulations were performed using HOOMD-blue (version 2.9.3) ^41^ with features extended using azplugins (version 0.10.1). ^42^ We simulated phase coexistence in rectangular simulation boxes (10 nm × 10 nm × 75 nm) with 500 chains initially placed in a dense slab with surface normals aligned with the *z*-axis. We also performed simulations of the same number of chains in cubic boxes of edge length 40 nm to study the formation of finite-sized aggregates.

## Results and Discussion

### Single-chain heteropolymer ensembles are matched by scaling attractions

Since our model sequences have the same degree of polymerization (*N* = 20), their single-chain dimensions (*i.e*., at infinite dilution) are mainly dictated by the fraction of polar monomers *X*_P_ and specific intramolecular interactions. Strong monomer attraction would lead to the collapse of the chain, mimicking poor solvent conditions, whereas pure repulsion would lead to chain expansion analogous to good solvent conditions. ^9,43^ Single-chain compactness has been shown to correlate with the propensity to phase separate both experimentally and computationally for a wide range of IDPs. ^44–46^ Here, we quantified single-chain compactness using the polymer’s radius of gyration *R*_g_, which we computed from the average of the trace of the gyration tensor.^38,47^ To facilitate comparison of our sequence- and model-dependent results, we used a purely hydrophobic polymer (*X*_P_ = 0) with *λ*_H_ = 1 and a purely hydrophilic polymer (*X*_P_ = 1) as reference sequences.

We first measured *R*_g_ for the different sequences and models while fixing *λ*_H_ = 1. As expected, we found that *R*_g_ increased with increasing *X*_P_, indicating a reduced hydrophobic character (Figs. 2a and S2a). Furthermore, the probability distribution of *R*_g_ widened with increasing *X*_P_, highlighting that the chains exhibited greater conformational fluctuations for large *X*_P_ (Fig. S2b). Since our primary goal is to study the effect of interaction model and sequence patterning on phase behavior, we tuned the monomer interactions so that all investigated polymers had a comparable single-chain *R*_g_ (and thus similar compaction at infinite dilution) as the purely hydrophobic reference sequence. To account for the decrease in the hydrophobicity of the chain with increasing *X*_P_, we initially attempted to rescale *λ*_H_ = 1/(1−*X*_P_) to maintain the same average hydropathy per monomer 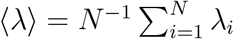 as the purely hydrophobic reference sequence. Scaling the interactions in this way did not, however, result in constant *R*_g_, highlighting the need for fine tuning of the interaction strengths for both models (Fig. S2a).

**Figure 2:**
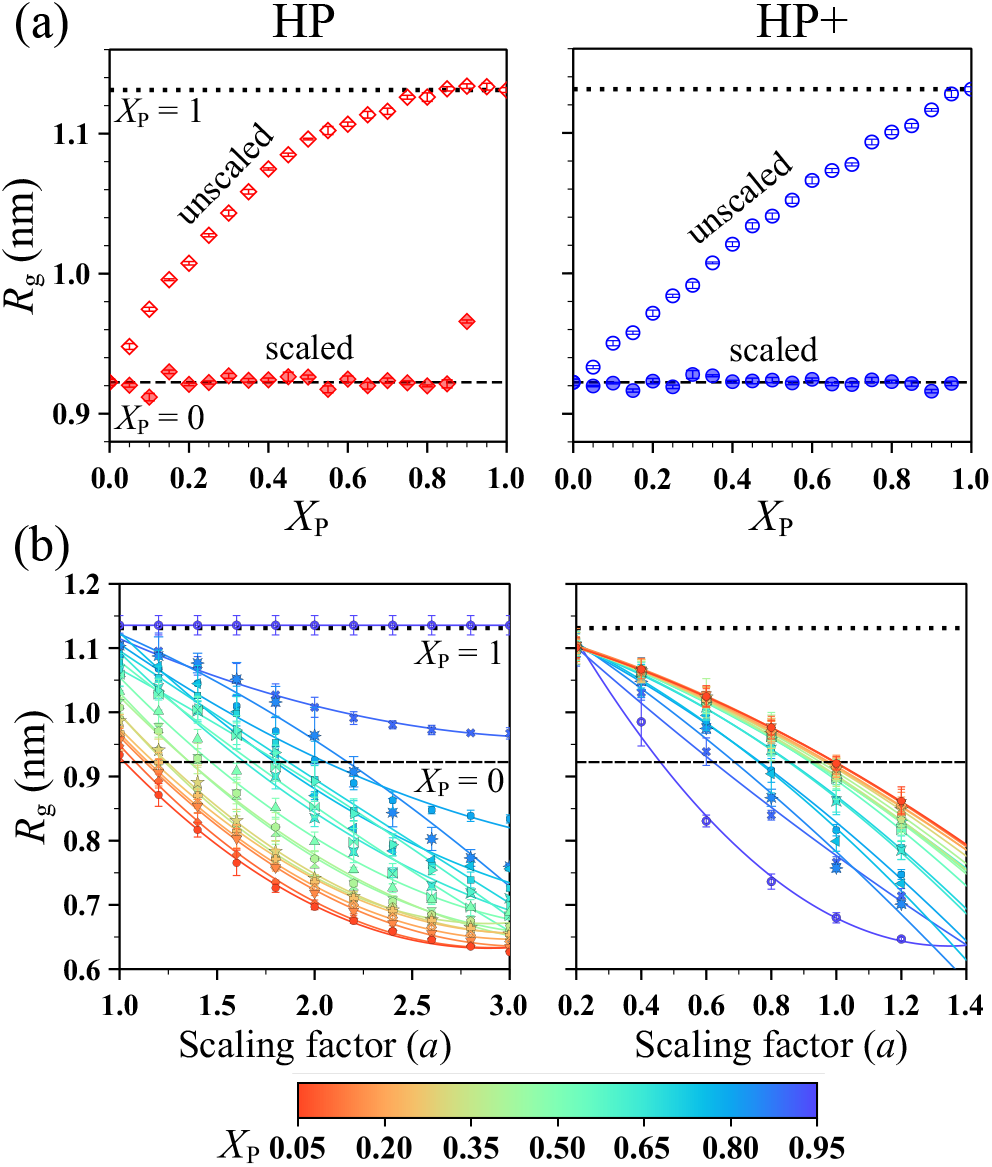
(a) Radius of gyration *R*_g_ as a function of *X*_P_ before scaling (*λ*_H_ = 1, open symbols) and after scaling (*λ*_H_ = *a*/(1−*X*_P_), closed symbols) for the HP (red) and HP+ (blue) models. (b) *R*_g_ as a function of scaling factor *a* using the HP and HP+ models. Results for purely hydrophobic (*X*_P_ = 0) and purely hydrophilic (*X*_P_ = 1) reference sequences are shown as dashed and dotted black lines, respectively.

Accordingly, we introduced an additional scaling factor *a* to modulate *λ*_H_ = *a*/(1 − *X*_P_). We selected *a* for each polymer by running a series of simulations to determine *R*_g_ as a function of *a* (Fig. 2b), then found the value of *a* that matched *R*_g_ to the hydrophobic reference sequence. We found *a*> 1 for the HP model, suggesting that the required monomer–monomer attractions are stronger compared to those needed to maintain the average hydropathy of the sequence (*i.e*., ⟨*λ*⟩ > 1). Conversely, we found *a*< 1 in the HP+ model and thus ⟨*λ*⟩ < 1. With these scaling factors applied to the HP model, we observed that for all values of *X*_P_, other than 0.90 and 0.95, *R*_g_ remained within 3% of that of the purely hydrophobic reference sequence (Fig. 2a). In the *X*_P_ = 0.90 sequence, strong attraction existed between the only two H monomers present in the sequence, and therefore their position within the sequence mainly dictated *R*_g_ (Figs. 2a and S3), while in the *X*_P_ = 0.95 case, the chain behaved as a purely hydrophilic chain due to the presence of only one H monomer. For the HP+ model, we matched *R*_g_ for all *X*_P_ including the 0.90 and 0.95 cases due to the additional attractive interaction between H and P monomers present in the model.

To further quantify the similarity established at the single-chain level, we next investigated if the match in average *R*_g_ also extended to the probability distributions *P* (*R*_g_). Indeed, we found similar *P* (*R*_g_) in the HP model for *X*_P_ ≲ 0.5, but the agreement quickly deteriorated for larger *X*_P_ (Fig. 3a), possibly due to the reduction in the number of available interaction sites (*i.e*., H monomers). For the HP+ model, we found a near perfect overlap of *P* (*R*_g_) (Fig. 3b), implying that the single-chain conformations in the HP+ model were almost identical to those of a purely hydrophobic reference sequence for all investigated *X*_P_ values. To further test this similarity, we considered the shape anisotropy ⟨*κ*^2^⟩, computed using the eigenvalues of the gyration tensor;^23,38^ the nonbonded potential energies *U*_nb_; and the relaxation time *τ*_e_ of the end-to-end vector autocorrelation function ^38^ (Fig. 3c). Remarkably, we found for almost all *X*_P_ values ⟨*κ*^2^⟩ ≈ 0.39, similar to the numerically determined value of a three-dimensional random walk,^48^ despite the very different sequence compositions. Again, we found that the HP+ model reproduced the reference state up to significantly larger *X*_P_ compared to the HP model. We observed similar trends in the nonbonded potential energy *U*_nb_: for the HP model, *U*_nb_ remained almost the same as in the purely hydrophobic reference sequence up to *X*_P_ ≈ 0.5, while in the HP+ model, it remained the same throughout the entire *X*_P_ range. Surprisingly, we did not see as pronounced a difference between the two models when considering the end-to-end vector relaxation times *τ*_e_ with varying *X*_P_. To check the sensitivity of the employed models and single-chain indicators, we considered three additional sequence sets in the range from *X*_P_ = 0 to *X*_P_ = 1 (RS1, RS2, and RS3 in Fig. S1) but did not find any strong sequence dependence (Figs. S3 and S4). Additionally, we found that the scaling factor *a* remained similar for all sequence sets considered in this work (Fig. S5).

**Figure 3:**
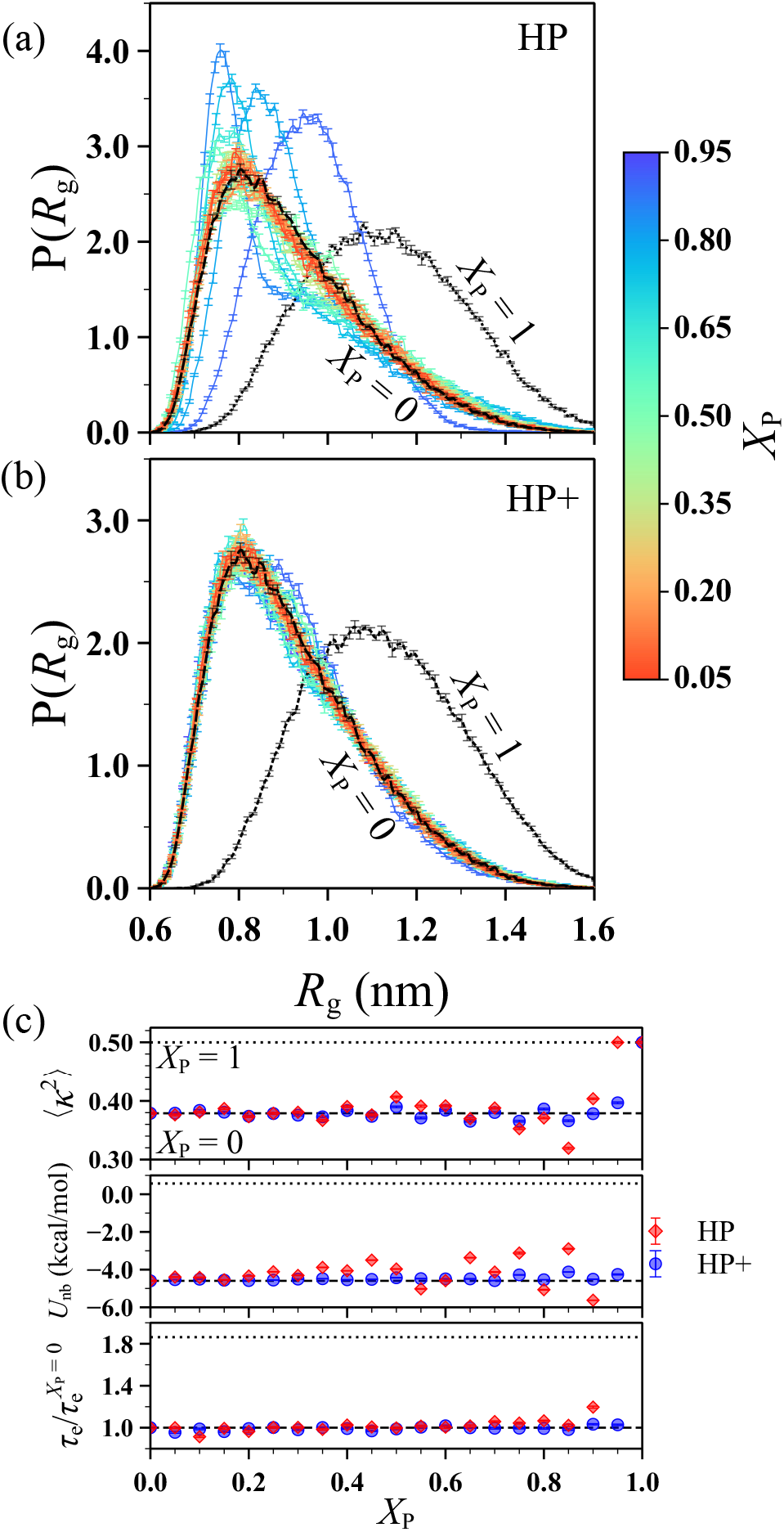
Probability distribution of radius of gyration *R*_g_ for the (a) HP and (b) HP+ models. (c) Shape anisotropy ⟨*κ*^2^⟩, nonbonded potential energy *U*_nb_, and end-to-end vector relaxation time *τ*_e_ as functions of *X*_P_ for the HP and HP+ models. The last quantity in (c) is normalized by that obtained for the purely hydrophobic reference sequence (*X*_P_ = 0). Results for purely hydrophobic and purely hydrophilic (*X*_P_ = 1) reference sequences are shown as dashed and dotted black lines, respectively.

To summarize, tuning the attraction strength between H monomers gave similar average single-chain properties and *P* (*R*_g_) as the purely hydrophobic reference sequence across for all sequences in the HP+ model, but only about half of the sequences in the HP model. The better match over the entire *X*_P_ range is due to the additional attractive interaction between H and P monomers in the HP+ model. The HP+ model is less sensitive to changes in the number of attractive monomers as compared to the HP model at the single-chain level and hence might be expected to better capture the condensed-phase behavior of the purely attractive homopolymer for different values of *X*_P_.

### Localized interactions reduce phase-separation propensity

Having established similar single-chain properties for both models, we next investigated the implications for the condensed-state properties of the sequences studied. According to homopolymer-based solution theories, intrachain contacts are translated to interchain contacts in concentrated solutions, thus linking the behavior at the single-chain level to phase-separation propensity.^9,46,49,50^ Panagiotopoulos and coworkers^51^ showed *via* on-lattice Monte Carlo simulations that in the limit of infinitely long homopolymer chains, the critical temperature, the Boyle temperature, and the coil–to–globule transition temperature are equivalent. Recent simulations of heteropolymeric model IDPs and naturally occurring IDPs have revealed a similar correlation between the extent of collapse at the single-chain level and phase-separation propensity.^46^ Extending the ideas above, corrections were proposed to account for the correlation between single-chain and condensed-state properties for charged heteropolymers.^14^ These prior works suggest that principles from homopolymer theory can be applied with reasonable accuracy when attractive monomers are distributed rather uniformly along the protein sequence. ^23^ With these considerations, we investigated whether the similarity established at the single-chain level by scaling monomer interactions also leads to similarities in phase behavior for our model sequences having strong localized or weak distributed interactions.

We performed direct coexistence simulations of the dilute and dense phases, computed time-averaged concentration profiles (Fig. 4), and extracted the coexistence concentrations of the dilute and dense phases (Fig. 5). In the HP model, we found similar dilute-phase concentrations *c*_sat_ for *X*_P_ ≤ 0.10, while beyond *X*_P_ = 0.15, we observed an increase in *c*_sat_ and a corresponding decrease in the phase-separation propensity (Figs. 4 and 5). We also found that the dense-phase concentration started to decrease beyond *X*_P_ = 0.15. For *X*_P_ ≥ 0.50, we found parabolic concentration profiles centered at the origin, indicative of the formation of clusters rather than a bulk condensed-phase (see representative simulation snapshots^52^ for *X*_P_ = 0.50 and *X*_P_ = 0.75 in Fig. 4). From these observations, we concluded that the threshold value above which the sequences did not phase separate is roughly 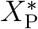 = 0.45 for the HP model.

**Figure 4:**
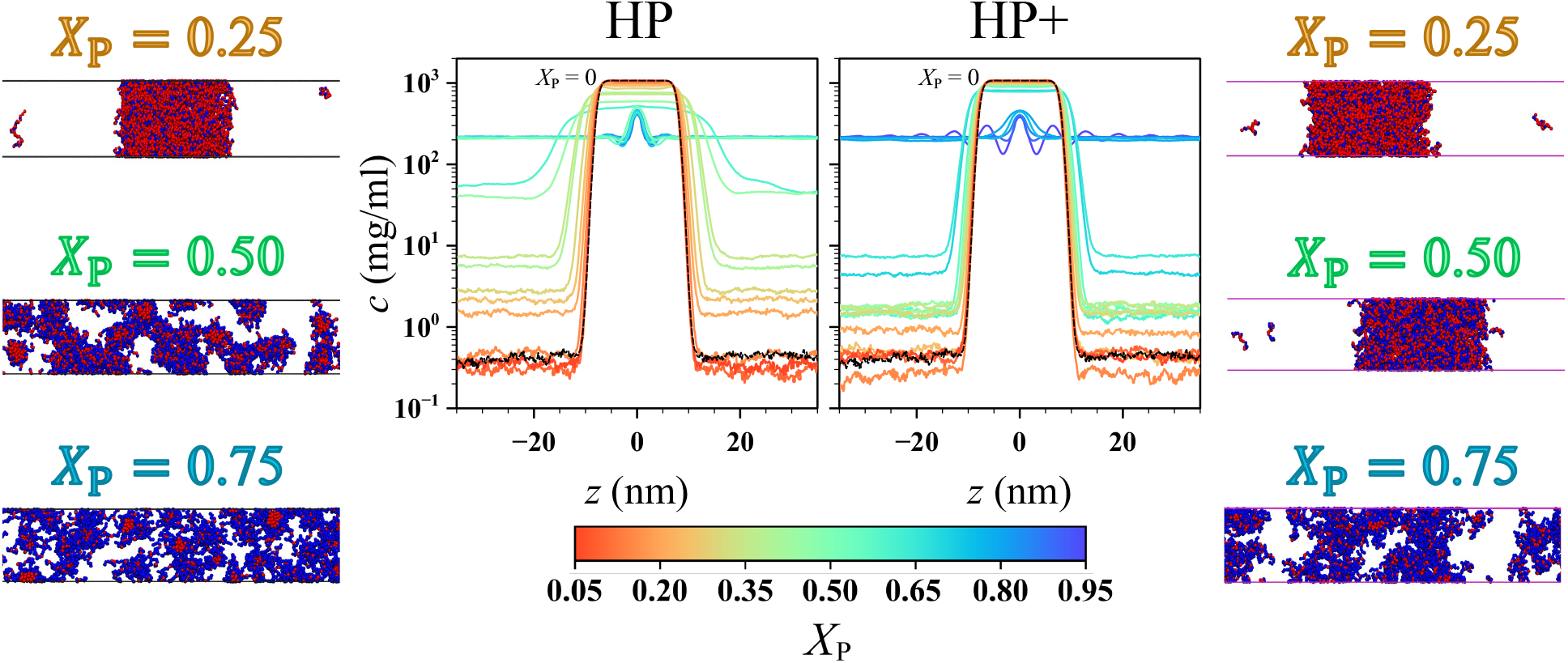
Concentration profiles estimated using direct coexistence simulations for the HP and HP+ models. The black line represents the purely hydrophobic reference sequence (*X*_P_ = 0). Representative simulation snapshots for *X*_P_ = 0.25, 0.50, and 0.75 are shown on the left (HP) and right (HP+) sides.

**Figure 5:**
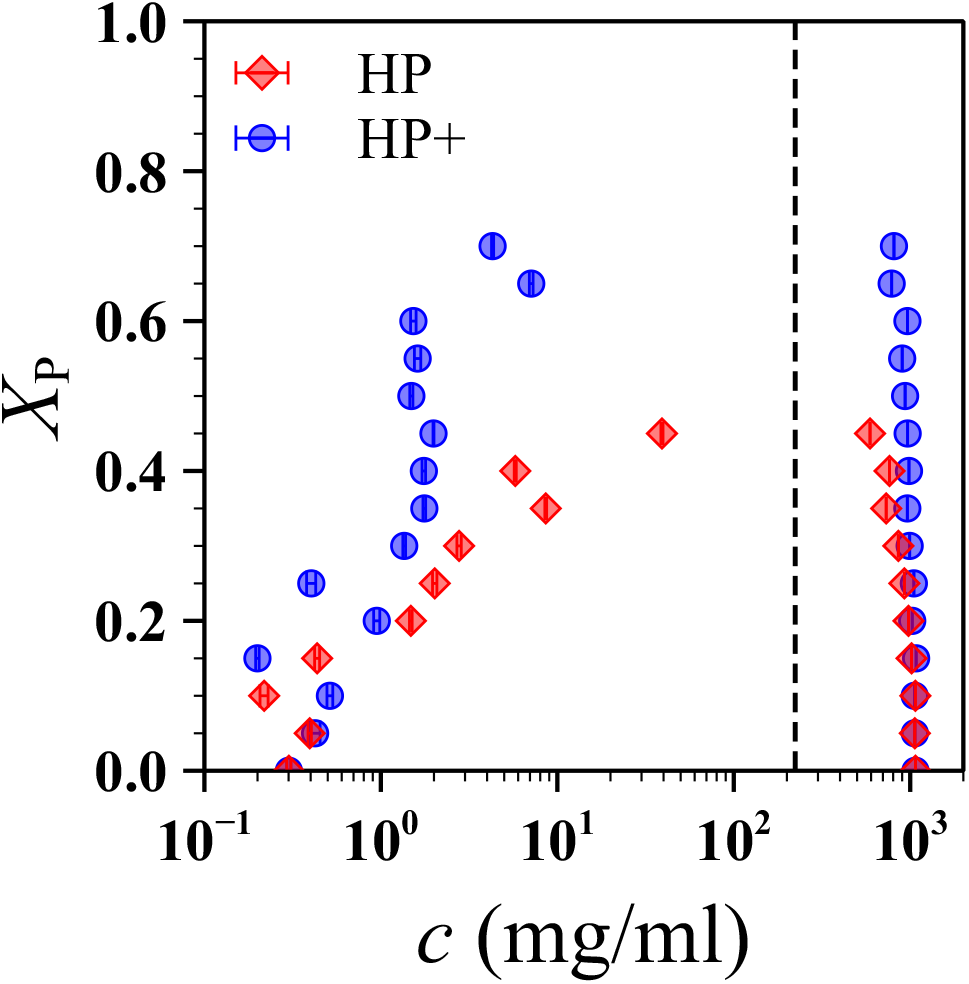
Phase diagrams for the HP and HP+ models. Concentrations of dilute and dense phases for all *X*_P_ values were extracted from the concentration profiles shown in Figure 4. Dashed line represents the total concentration of the coexistence simulation.

In the HP+ model, phase separation persisted for higher *X*_P_ compared to the HP model (Figs. 4 and 5), highlighting the stabilizing role played by the cross interactions between H and P monomers. We found that the coexistence concentrations remained almost the same only for *X*_P_ ≤ 0.15. Beyond *X*_P_ = 0.20, *c*_sat_ increased, indicative of lower phase-separation propensity as compared to the purely hydrophobic reference sequence under the conditions of this study. However, compared to the HP model, we observed cluster formation only for large *X*_P_ values (*X*_P_ ≥ 0.75; Fig. 4). Based on this finding, we defined 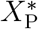 = 0.70 for the HP+ model, which suggests that at least 30% of the sequence must consist of hydrophobic monomers to form a liquid-like condensate at the simulated temperature. We find that for both models, matching the single-chain dimensions through scaling of interactions leads to a dramatic increase in the threshold values (0.1 to 0.45 for HP and 0.25 to 0.70 for HP+) for phase separation when compared to the unscaled interactions (Fig. S6). This finding indicates that scaling the interactions between H monomers in order to match the single-chain dimensions did offset the effect of changes in composition in both models. However, the extent to which this similarity established at the single-chain level could compensate for changes in composition of the sequence was limited.

The sequences with *X*_P_ around the threshold 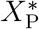 value (*X*_P_ = 0.35 and 0.40 in the HP model, and *X*_P_ = 0.65 and 0.70 in the HP+ model) formed a wider dense phase slab with its concentration about 75% of that of the purely hydrophobic reference sequence (*X*_P_ = 0). To investigate whether this widening is due to the decrease in concentration or the formation of void volume within the slab, we computed the radial distribution function *g*(*r*) between H monomers for these sequences (Fig. S7). We found only a marginal increase in the magnitude of the first peak in *g*(*r*) compared to the *X*_P_ = 0 sequence, highlighting the absence of preferential patterning of H and P monomers in the dense phase of these sequences. Both H and P monomers were well mixed within the dense phase of all sequences up to 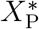 (Fig. 5), further substantiating their ability to form homogeneous liquid-like condensates.

We next probed whether the system could form a condensed-phase above the threshold 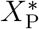 values (0.45 for the HP model and 0.70 for the HP+ model) by further increasing the attraction between H monomers. To this end, we ran simulations for all cases that did not form stable slabs using the HP+ model with *a* increased to 2, 5, and 10 times that needed to match the single-chain *R*_g_. At such high interaction strengths, we observed the formation of highly patterned slabs as reflected in their concentration profiles (Fig. S8). Additionally, the concentrations of the dense phase of these sequences showed a system size dependence, analogous to those classified as aggregates by Panagiotopoulos and coworkers. ^53^ From these observations, we concluded that insufficient interaction strength was likely not the reason why the sequences above 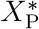 did not form liquid-like condensates.

To investigate the sequence dependence of our observations, we also characterized the phase behavior of three additional sequence sets (RS1, RS2, and RS3 in Fig. S1). We found that the threshold value 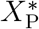 showed a stronger sequence dependence than the single-chain behavior (Fig. S9). However, 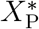 for the HP+ model was always higher than that observed for the HP model for all sequence sets, consistent with our primary sequence set. Taken together, we found that matching single-chain properties did not prevent the decrease in the phase-separation propensity with increasing localization of interactions within the sequences, indicating that interaction strength alone did not dictate the formation of a homogeneous condensed-phase but rather a combination of interaction strength and the number of attractive monomers.

### Scaled interactions lead to favorable interchain interactions for all sequence compositions

A possible reason for the reduction in phase-separation propensity with increasing *X*_P_ could be the decreasing favorability of interchain interactions despite favorable intrachain interactions. To quantify the strength of interaction between a pair of chains, we determined the chain–chain second virial coefficient *B*_22_. Positive *B*_22_ indicates effective repulsion, while negative *B*_22_ indicates effective attraction. We used a combination of replica exchange Monte Carlo and umbrella sampling methods to compute the dilute chain–chain potential of mean force (PMF), *w*(*r*). Then, *B*_22_ was computed as^32,46^

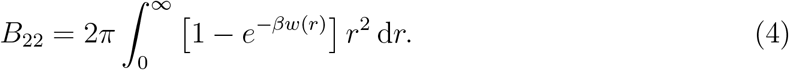

The above method is known to estimate *B*_22_ with a high degree of statistical certainty.^32,46^ Fig. 6a shows the PMF for select *X*_P_ values (see Figs. S10 and S11 for the entire set), while the resulting *B*_22_ values for all *X*_P_ values are shown in Fig. 6b. For *X*_P_ ≲ 0.40, both models showed highly similar PMF to that of the purely hydrophobic reference sequence in terms of both the location and depth of the attractive well. At intermediate to high values of *X*_P_, we observed changes between the two models: PMFs of the HP+ model were shallower and closer to the purely hydrophobic reference sequence as compared to the HP model (Fig. 6a). This behavior implies that the effective chain-chain interactions in the HP model are stronger than in the HP+ model. For *X*_P_ = 0.90, the PMF obtained for the HP model is an order of magnitude more attractive than that of the HP+ model, possibly owing to the higher attraction strength needed to match the single chain dimensions using only two attractive H monomers.

**Figure 6:**
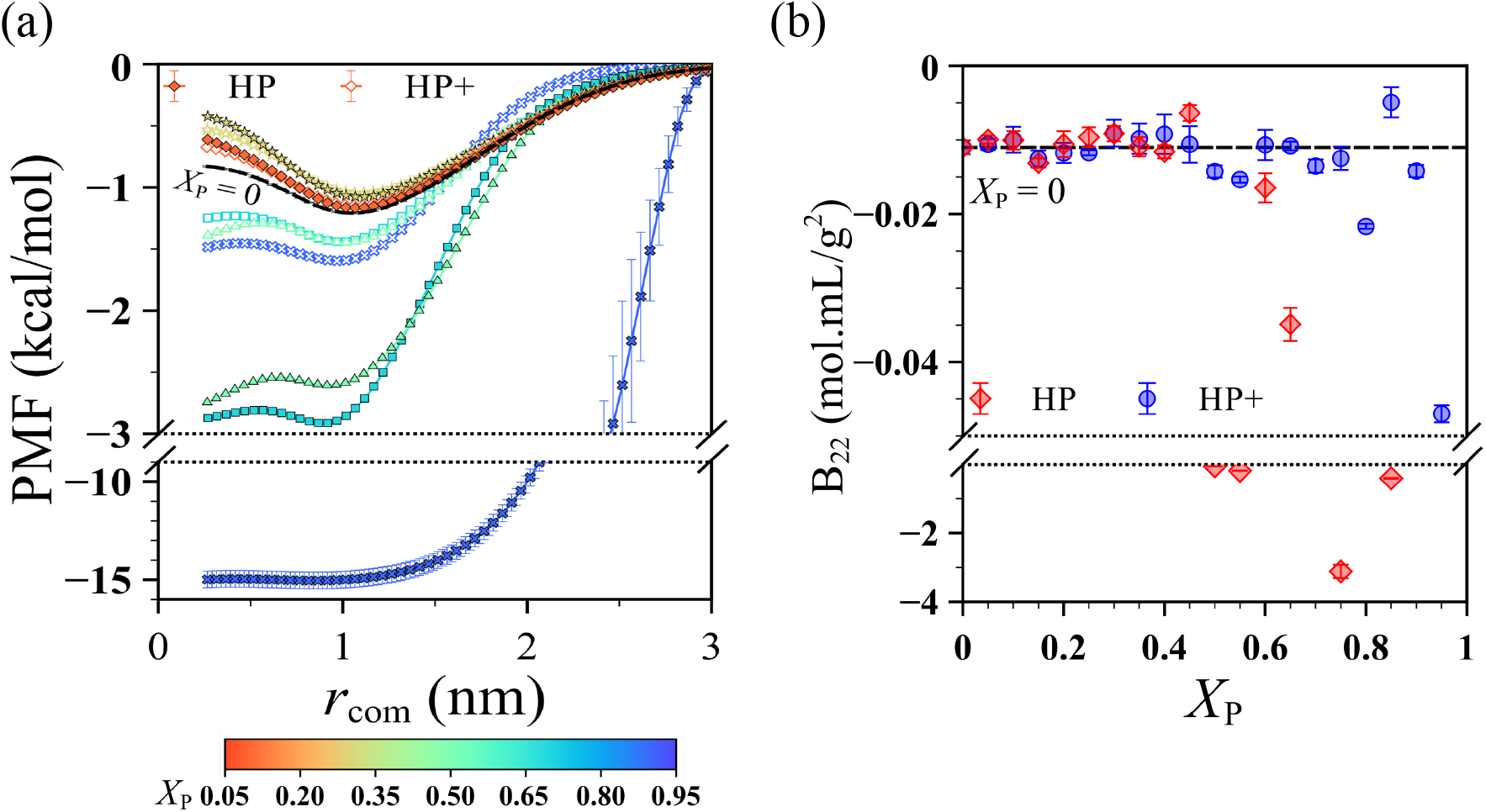
(a) Potential of mean force (PMF) as a function of center-of-mass distance *r*_com_ between two chains for select *X*_P_ values using the HP (filled symbols) and HP+ (open symbols) models. (b) Second viral coefficient *B*_22_ obtained from the PMFs for the HP and HP+ models.

This comparison shows that the similar behavior achieved at the single-chain level for all values of *X*_P_ translates to similar interactions at the two-chain level, with slightly negative *B*_22_ values up to roughly 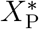. Beyond 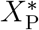, for both models, we see that the *B*_22_ values fluctuate, with the HP model showing more negative values than the HP+ model indicating that the effective attraction between chains is stronger in the HP model than the HP+ model as the number of attractive sites decreases. In some cases (*X*_P_ ≥ 0.50 for the HP model and *X*_P_ = 0.80, 0.90, and 0.95 for the HP+ model), the chain-chain interactions are more favorable compared to the purely hydrophobic case, which implies that from a pairwise interaction standpoint, these cases would be more prone to chain–chain association than the purely hydrophobic reference sequence. However, we did not observe the formation of a homogeneous condensate in those cases. This discrepancy prompts the question: what limits the system from undergoing phase separation if interactions remain favorable?

### Beyond the threshold for phase separation, finite-sized clusters are formed

Motivated by our finding that sequences with *X*_P_ >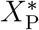
do not form a condensed homogeneous phase despite exhibiting strong chain-chain attraction, we next investigated the energetically favorable assemblies formed by such sequences using both models. To this end, we let the chains self-assemble within a cubic simulation box, and then performed a clustering analysis of the resulting aggregates. Specifically, we considered two chains to be part of the same cluster if the distance between monomers of two chains was less than 1.5*σ*, and then computed the probability of finding a chain in a cluster of size *N*_c_ (Fig. 7). The probability distributions *P* (*N*_c_) were relatively insensitive to the distance criteria chosen for analysis (Fig. S12). To rule out possible finite-size effects resulting from different box sizes and geometries (slab vs. cubic), ^46,54^ we first performed the analysis on sequences that undergo phase separation (*i.e*., *X*_P_ < 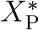; Fig. S13). Consistent with the results from slab simulations, we found that the chains predominantly resided in a single large cluster for these sequences.

Beyond 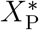, the probability of finding chains within smaller clusters was nonzero for both models (Fig. 7), indicating that interchain interactions are still favored in the limit of multiple chains. Interchain contacts are more favored in the case of the HP model as compared to the HP+ model, reflected by the higher probability of observing clusters of chains as compared to single chains in solution. The attraction between H and P monomers in the HP+ model allowed isolated chains to form intrachain contacts for all but *X*_P_ = 0.95, which formed micelles with their cores consisting of H monomers from multiple chains (Fig. 7). We hypothesized that the different behavior exhibited for *X*_P_ = 0.95 in the HP+ model could be due to the rather uniform patterning of H monomers. Indeed, in the case of a highly patterned sequence set (RS4 in Fig. S1), we found that all systems beyond 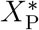 preferably formed small micellar clusters, thus highlighting the role of the arrangement of H monomers in dictating the formation of interchain or intrachain contacts (Fig. S14).^30^ Despite some differences between the specific cluster morphologies in the two models, it is clear that the general emergence of smaller clusters beyond 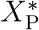 is model-independent. This finding further highlights that the formation of a single high concentration phase is governed by the interplay between favorability of interactions and the number of available interaction sites. ^55–58^ Beyond 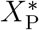, the system is limited by the number of available interaction sites (or valence), resulting in its inability to form a single continuous stable condensed-phase.

**Figure 7:**
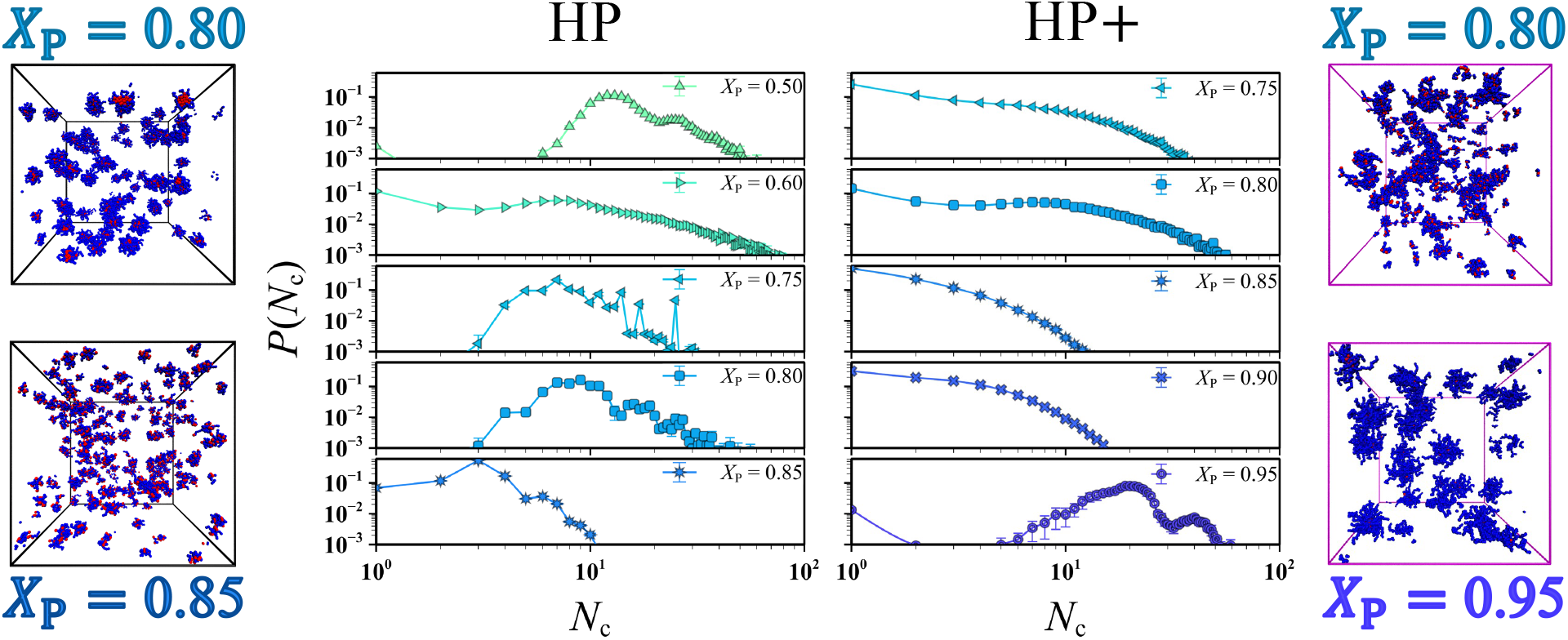
Probability of finding a chain in a cluster of size *N*_c_ for select *X*_P_ values that did not undergo phase separation in the HP and HP+ models. Representative simulation snapshots for *X*_P_ = 0.80 and 0.85 on the left side correspond to the HP model, while those for *X*_P_ = 0.80 and 0.95 on the right side correspond to the HP+ model.

## Conclusions

In this computational study, we comprehensively investigated the interplay between the localization and strength of interactions in dictating the phase behavior of model disordered proteins consisting of hydrophobic (H) and polar (P) monomers. We systematically varied the fraction of P monomers in the sequences, *X*_P_, and considered two model classes, representing interaction scenarios with either strong localized or weak distributed attractions. To establish a common ground for the large variation in IDP sequences, we carefully scaled the attraction strength in each case to match the single-chain dimensions of an attractive homopolymer, thereby providing all of the model proteins an equal opportunity to phase separate, from a single-chain perspective. To our surprise, scaling the interactions this way also led to similar conformational ensembles, nonbonded potential energies, and chain-level dynamics in both models for almost all investigated sequences. Based on the expectation that intramolecular interactions will translate directly to the intermolecular level, we probed the propensity of the IDPs to phase separate. We found similar phase-separation propensity for IDPs with sufficiently low *X*_P_, where a significant scale up in the attraction strength counterbalanced the decreasing number of H monomers. As *X*_P_ increased, however, the phase-separation propensity dramatically declined because of limited number of attractive monomers. Beyond a certain *X*_P_, the model IDPs did not form a continuous condensed-phase anymore, but instead self-assembled into finite-sized aggregates. These deviations from the reference attractive homopolymer were much more pronounced for the model with strong localized interactions, suggesting that weak distributed interactions between multiple residue types may better stabilize the liquid-like condensed-phase of disordered proteins.^10,18,24,31^ We believe that our work, performed at multiple scales, will aid in the mechanistic understanding of biomolecular phase separation and in the development of theoretical models to accurately capture the interactions of disordered proteins.

## Acknowledgments

This material is based on the work supported by the National Institute of General Medical Science of the National Institutes of Health under the grant R01GM136917 and the Welch Foundation under the grant A-2113-20220331. AN acknowledges funding by the Deutsche Forschungsgemeinschaft (DFG, German Research Foundation) through project 470113688. Y.C.K. is supported by the Office of Naval Research through the U.S. Naval Research Laboratory base program. Further, we acknowledge the Texas A&M High Performance Research Computing (HPRC) for providing the computational resources required to complete this work.

