## Supplementary Material for "Role of Strong Localized vs. Weak Distributed Interactions in Disordered Protein Phase Separation"

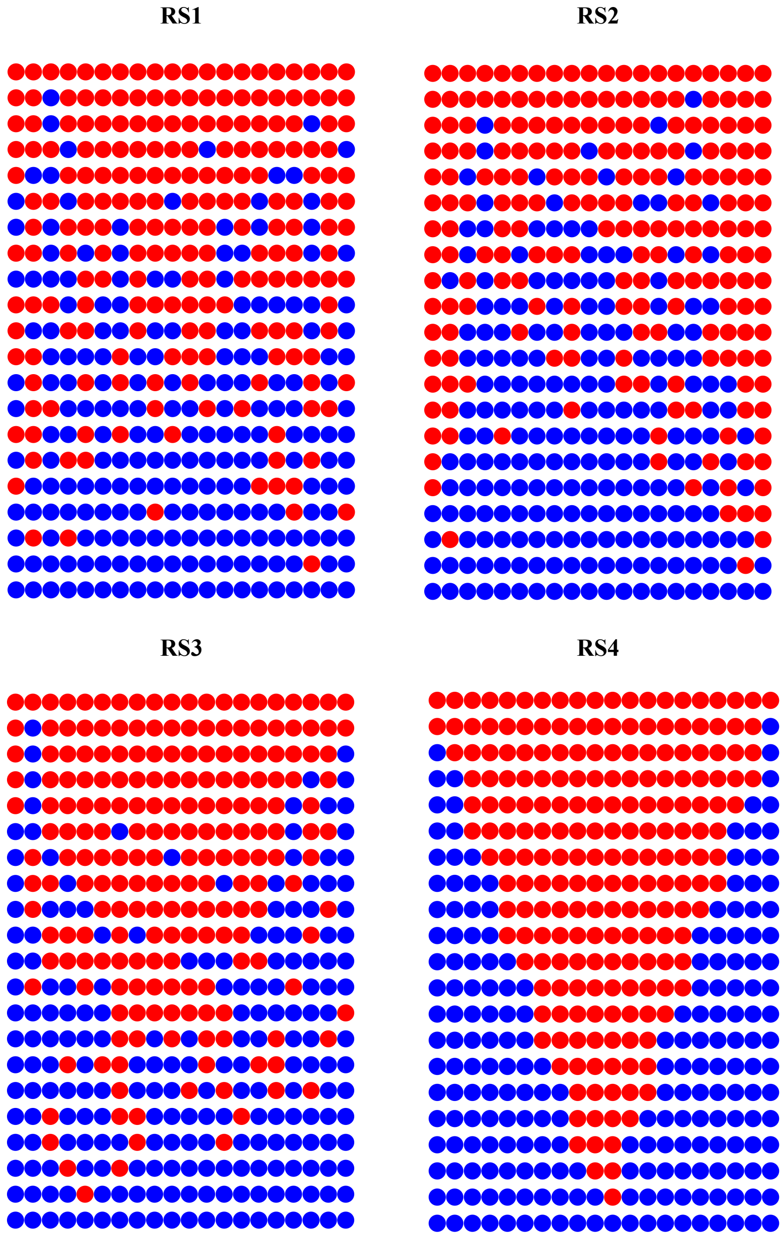


Figure S1: Three randomly generated sequence sets (RS1, RS2, and RS3) as well as a highly patterned sequence set (RS4). Red and blue beads correspond to hydrophobic (H) and polar (P) monomers, respectively.

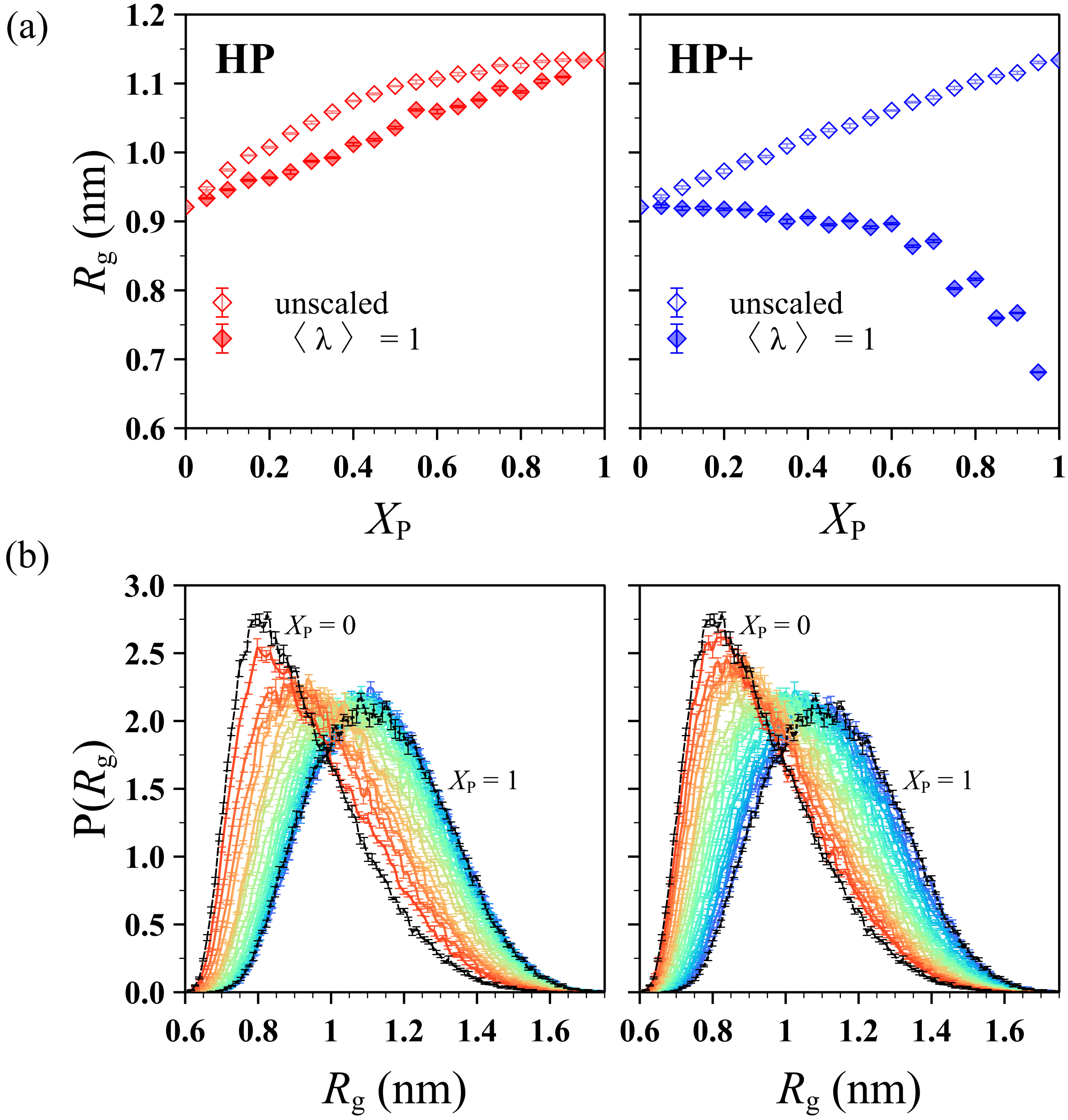


Figure S2: (a) Radius of gyration $R_g$ as a function of $X_P$ before scaling ($\lambda_H = 1$, open symbols) and after scaling ($\lambda_H = 1/(1 - X_P)$ that maintains average hydropathy per monomer $\langle\lambda\rangle$ of the chain at 1, closed symbols) for the HP (red) and HP+ (blue) models. (b) Probability distribution of $R_g$ when $\lambda_H = 1$ for the HP (left) and HP+ (right) models. The line color, ranging from red to purple, indicates increasing $X_P$. The distributions for purely hydrophobic ($X_P = 0$) and purely hydrophilic ($X_P = 1$) chains are shown as dashed and dotted black lines, respectively.

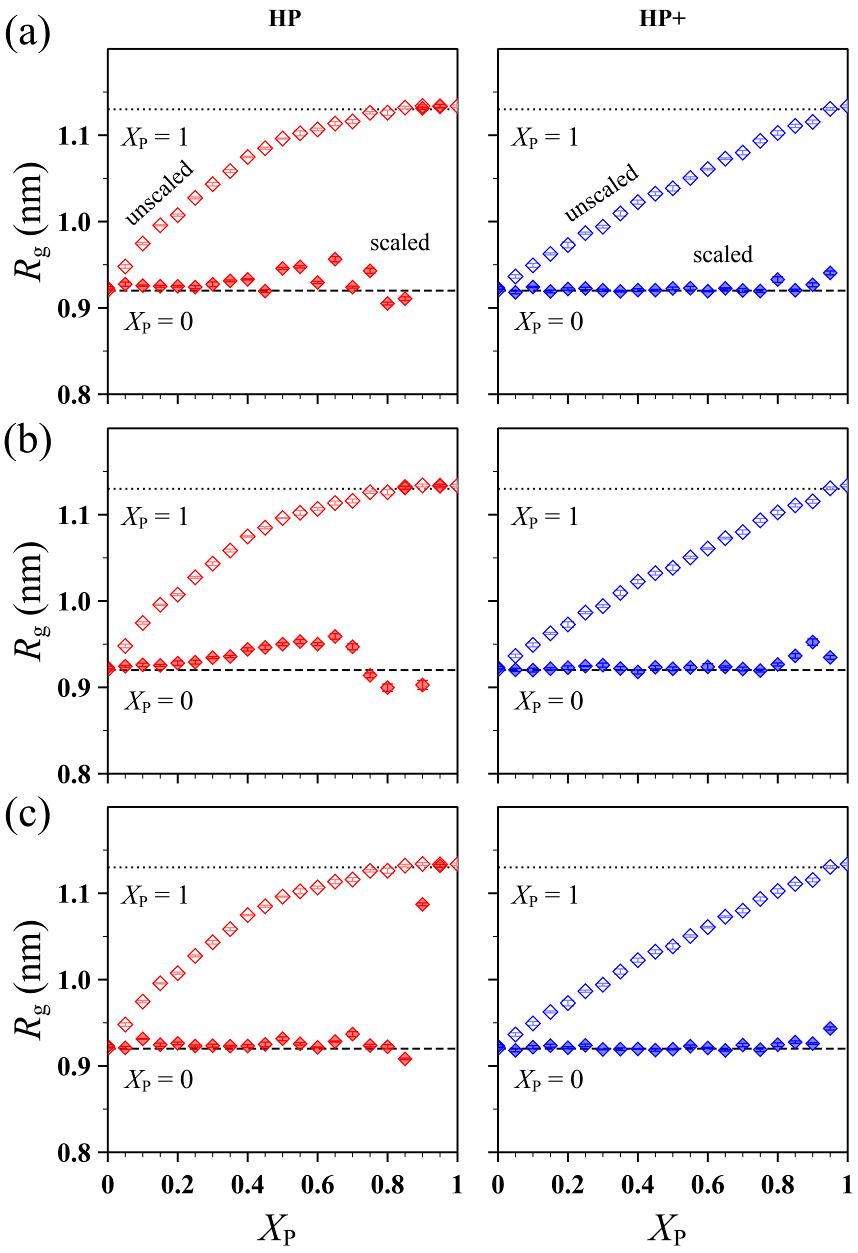


Figure S3: Radius of gyration $R_g$ as a function of $X_P$ before scaling ($\lambda_H = 1$, open symbols) and after scaling ($\lambda_H = a/(1 - X_P)$, closed symbols) for the HP (red) and HP+ (blue) models: (a) RS1, (b) RS2, and (c) RS3. Results for purely hydrophobic ($X_P = 0$) and purely hydrophilic ($X_P = 1$) chains are shown as dashed and dotted black lines, respectively, in all subplots.

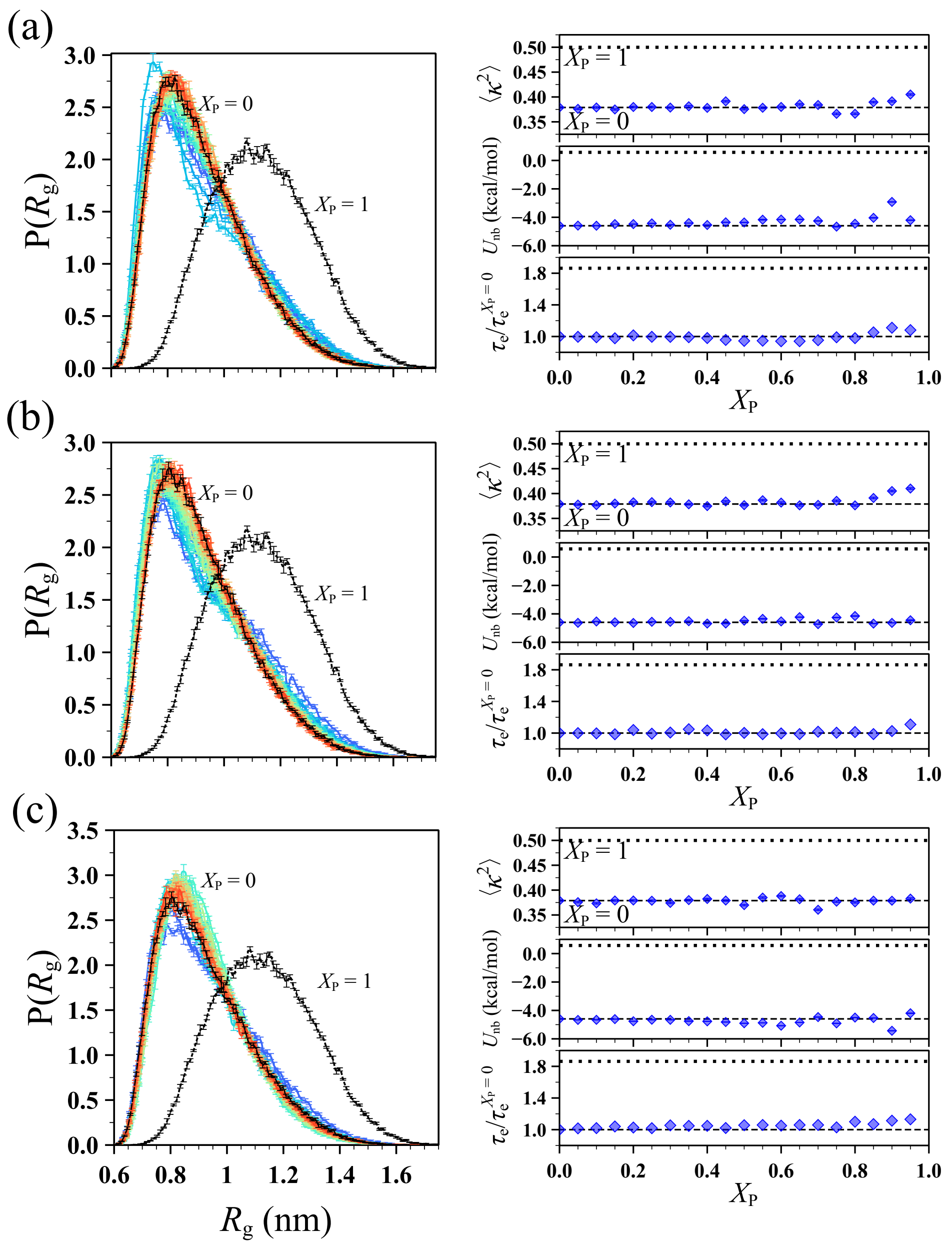


Figure S4: Probability distribution of radius of gyration $R_g$ (left) and plots showing shape factor $\langle\kappa^2\rangle$, nonbonded potential energy $U_{nb}$, and end-to-end vector relaxation time $\tau_e$ as functions of $X_P$ (right) for the HP+ model: (a) RS1, (b) RS2, and (c) RS3. In $P(R_g)$ plots, the line color, ranging from red to purple, indicates increasing $X_P$. The values of $\tau_e$ are normalized by that obtained for the purely hydrophobic chain ($X_P = 0$). Results for purely hydrophobic ($X_P = 0$) and purely hydrophilic ($X_P = 1$) chains are shown as dashed and dotted black lines, respectively, in all subplots.

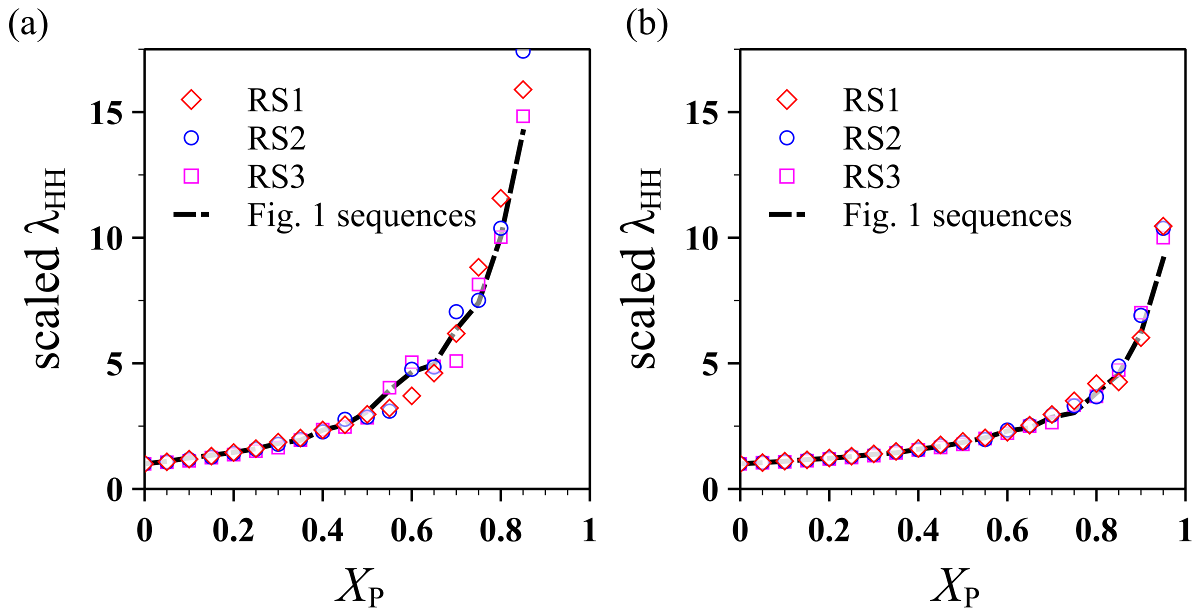


Figure S5: Comparison of scaled $\lambda_{HH}$ [$\lambda_{HH} = \lambda_H = a/(1 - X_P)$] between the three randomly generated sequence sets (RS1, RS2, and RS3) as well as the primary sequence set shown in Fig. 1 for the (a) HP and (b) HP+ models.

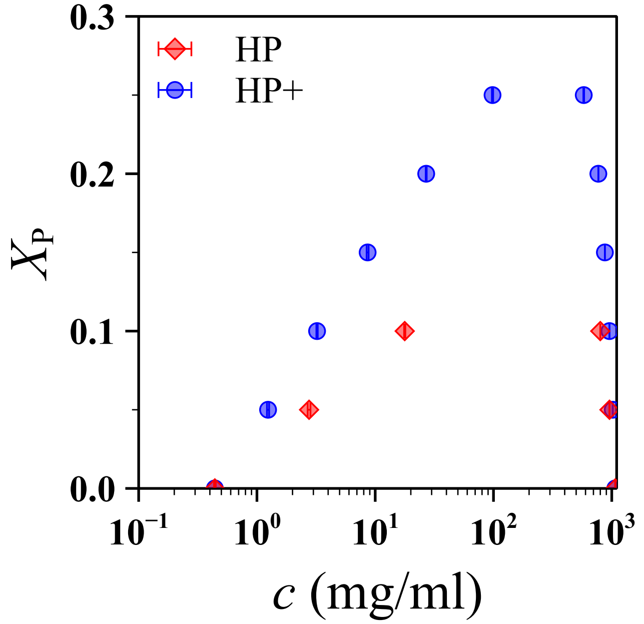


Figure S6: Phase diagrams for the HP and HP+ models when interactions are fixed at $\lambda_H = 1$. Concentrations of dilute and dense phases for all $X_P$ values were extracted from the corresponding concentration profiles.

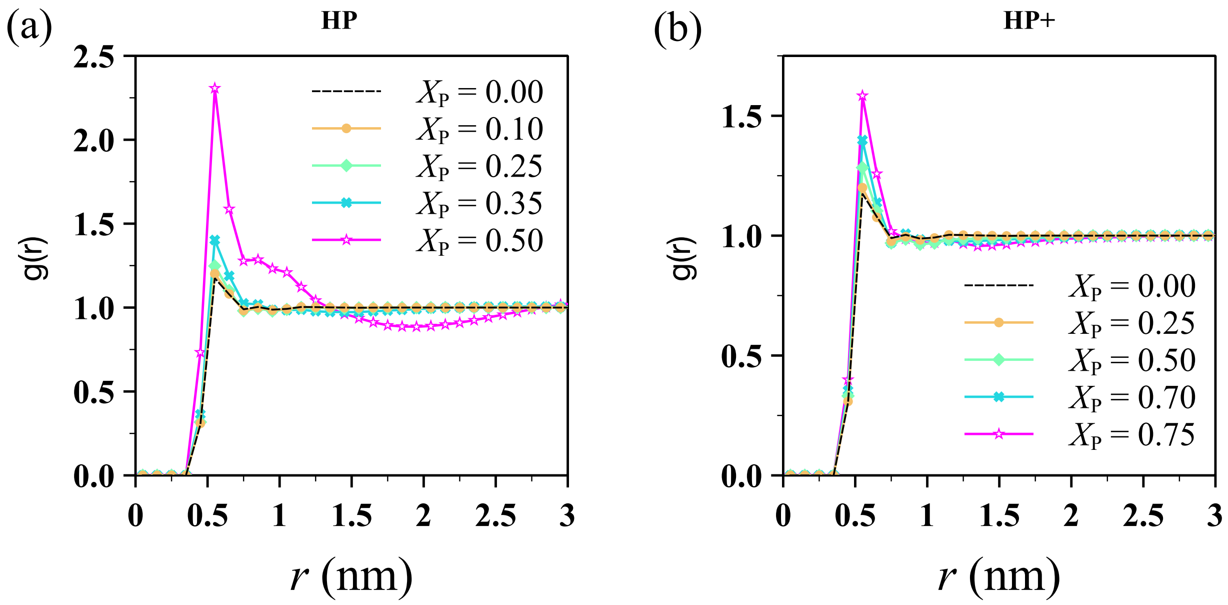


Figure S7: Radial distribution function $g(r)$ for different sequences that phase separated in the (a) HP model ($X_{\mathrm{P}} \leq X_{\mathrm{P}}^{*} = 0.45$) and (b) HP+ model ($X_{\mathrm{P}} \leq X_{\mathrm{P}}^{*} = 0.70$). The sequences that did not phase separate whose $X_{\mathrm{P}}$ is immediately above the threshold $X_{\mathrm{P}}^{*}$ (*i.e.*, $X_{\mathrm{P}} = 0.50$ in the HP model and $X_{\mathrm{P}} = 0.75$ in the HP+ model) are also shown for comparison.

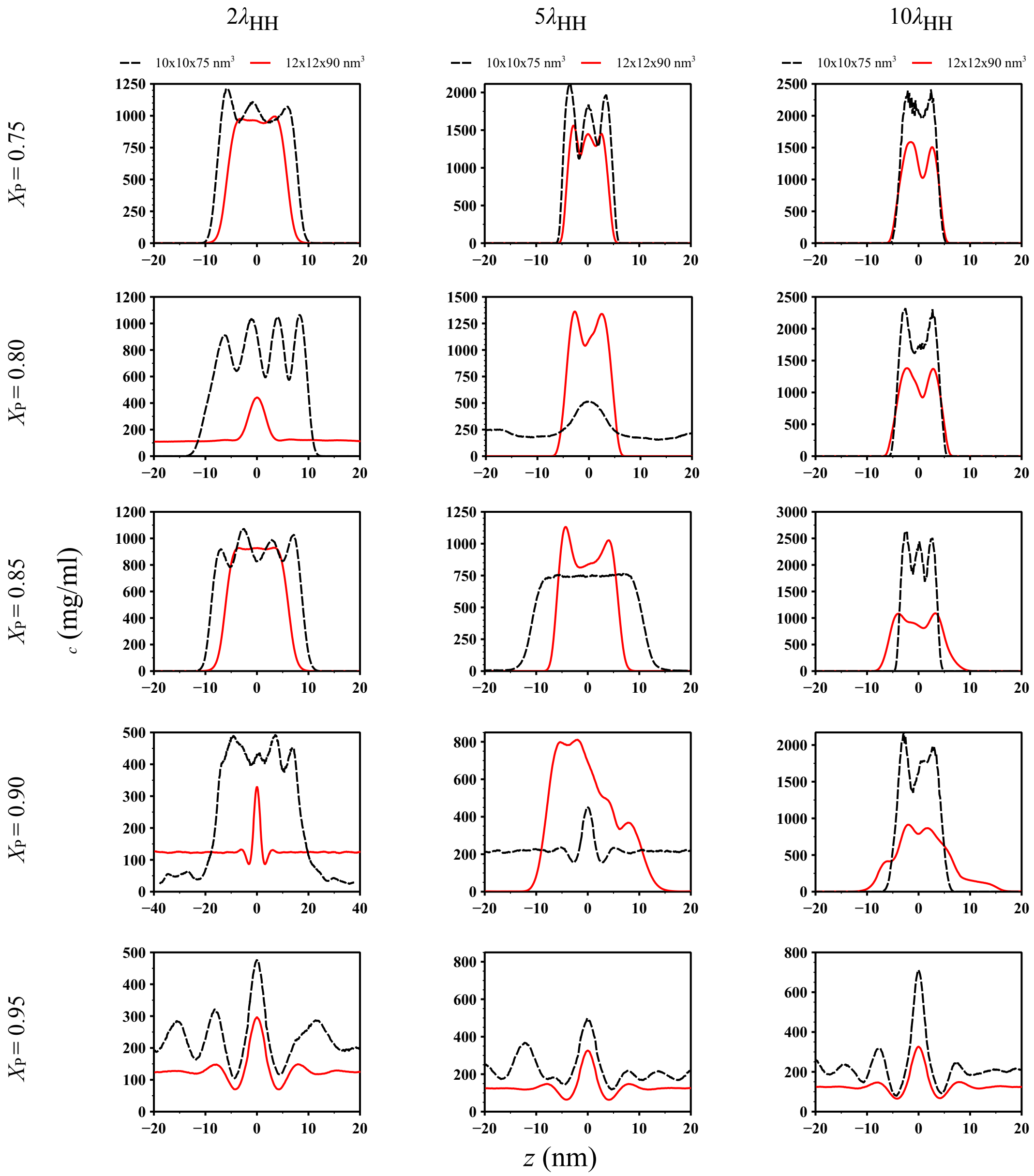


Figure S8: Concentration profiles for $X_{\mathrm{P}}$ values that did not undergo phase separation (*i.e.*, $X_{\mathrm{P}} > X_{\mathrm{P}}^{*} = 0.70$) in the HP+ model at $2, 5,$ and $10$ times the interaction strength $\lambda_{\mathrm{H}}$ required to match the radius of gyration $R_{\mathrm{g}}$ to that of the purely hydrophobic chain in a rectangular box of two different sizes: $10\,\mathrm{nm} \times 10\,\mathrm{nm} \times 75\,\mathrm{nm}$ (dashed black lines) and $12\,\mathrm{nm} \times 12\,\mathrm{nm} \times 90\,\mathrm{nm}$ (solid red lines).

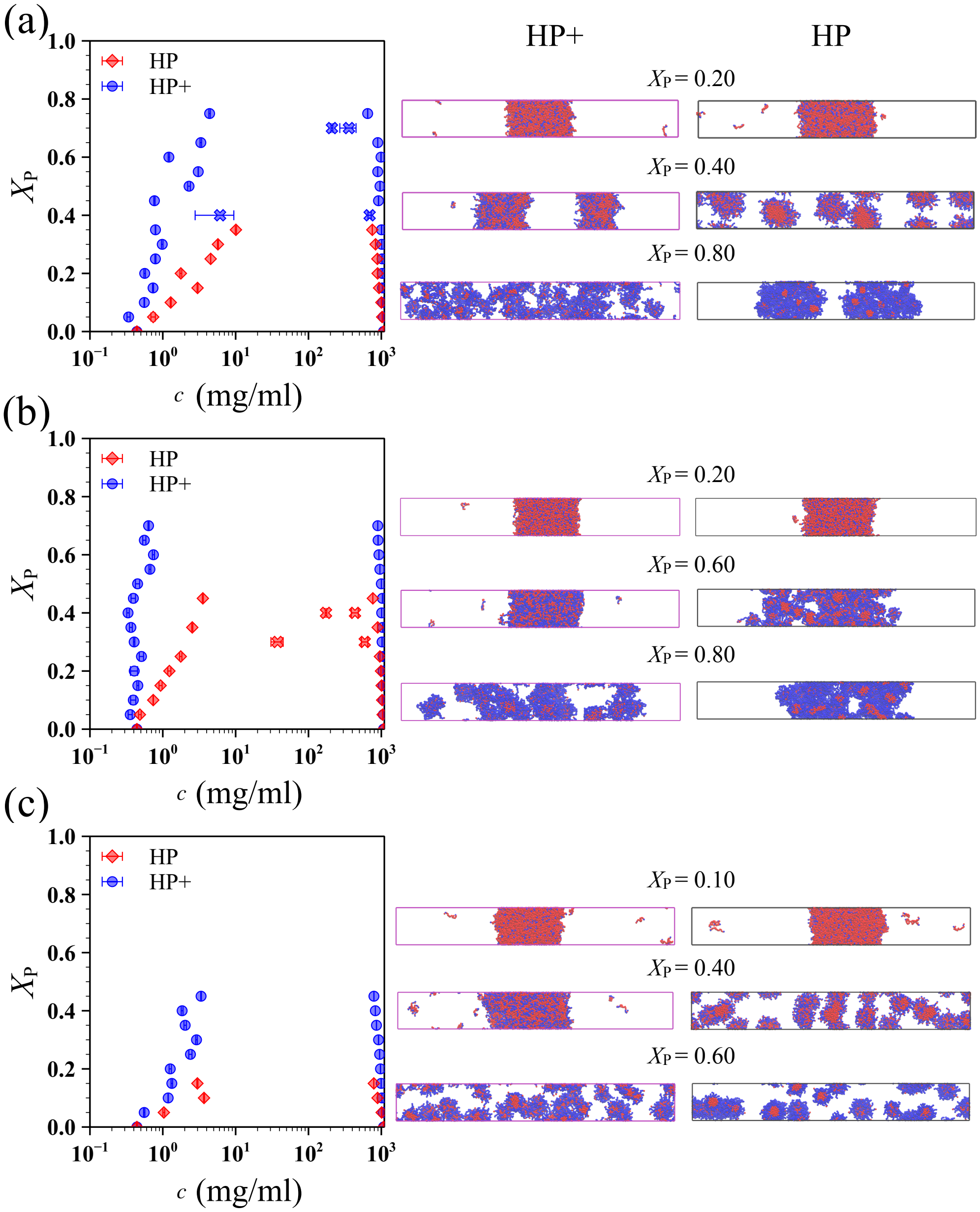


Figure S9: Phase diagrams and representative simulation snapshots for select $X_P$ values in the HP and HP+ models: (a) RS1, (b) RS2, and (c) RS3. Data points shown as crosses for RS2 correspond to $X_P$ values that did not phase separate even though they fell below the identified threshold $X_P^*$ value.

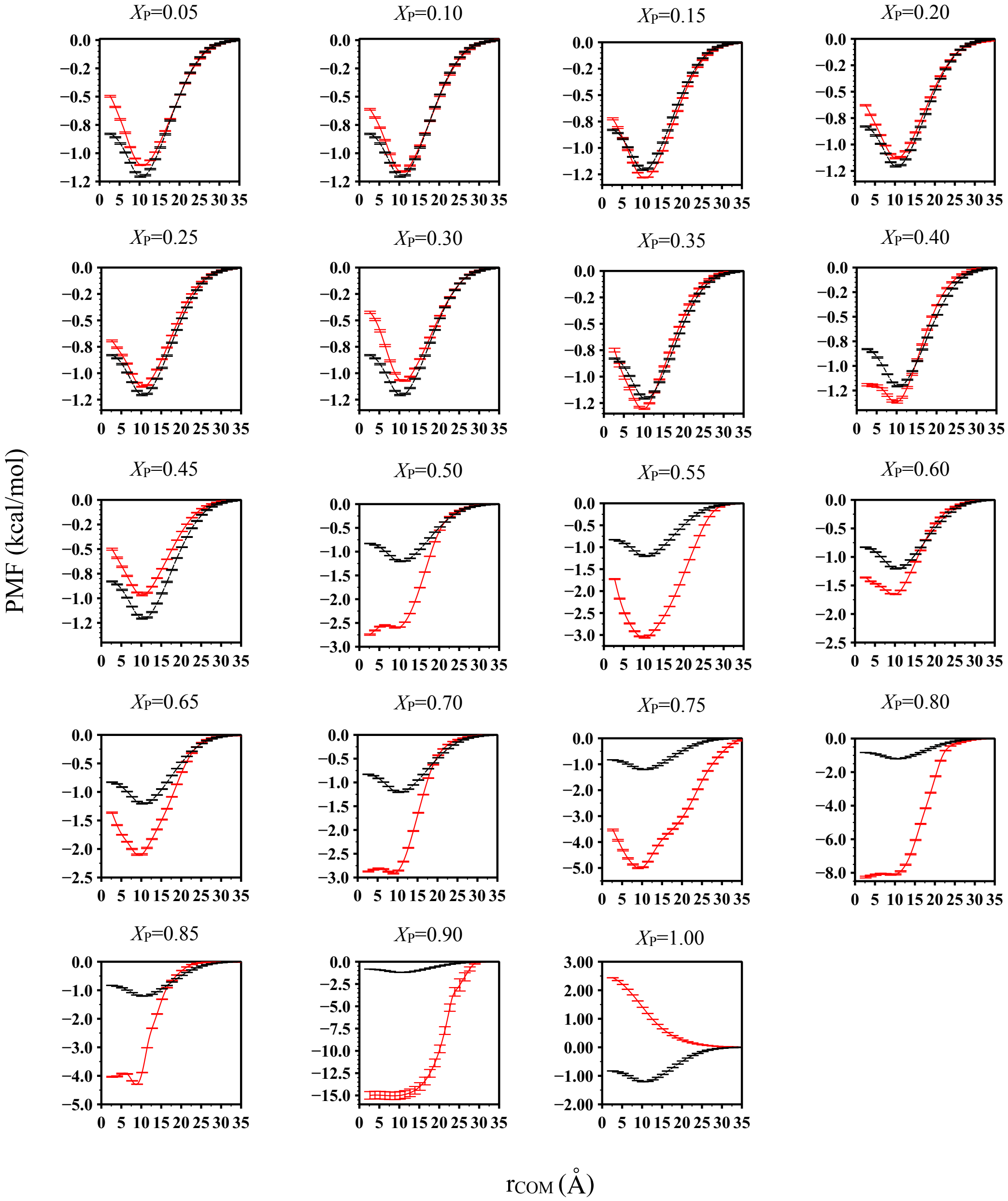


Figure S10: Potential of mean force (PMF) for all $X_{\mathrm{P}}$ values in the HP model. The PMF of the purely hydrophobic sequence ($X_{\mathrm{P}} = 0$; black dashed line) is shown as a reference in all subplots. Scaling factor $a$ to match the radius of gyration $R_{\mathrm{g}}$ for $X_{\mathrm{P}} = 0.95$ cannot be obtained and hence, its PMF is not shown in the plot.

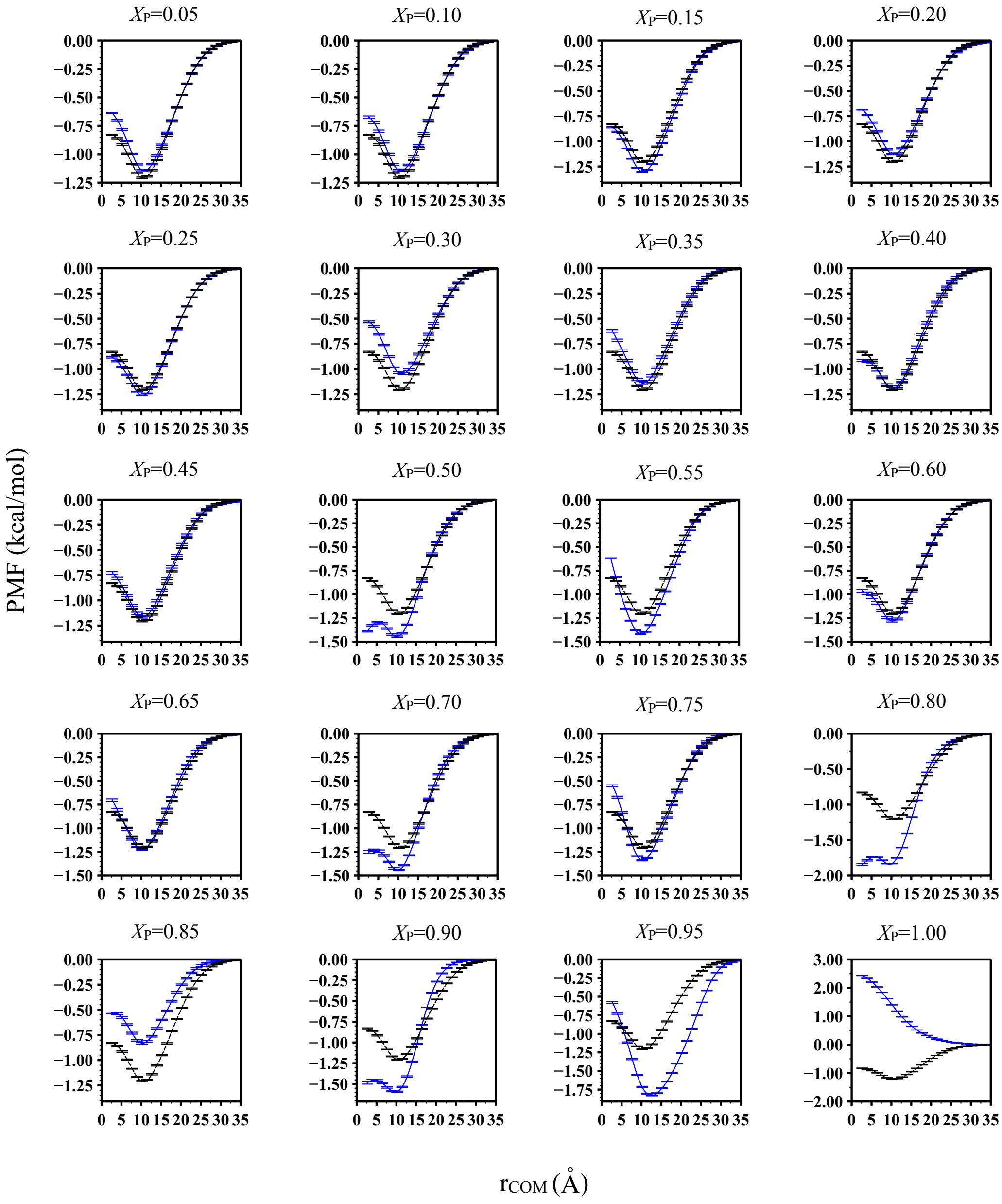


Figure S11: Potential of mean force (PMF) for all $X_{\mathrm{P}}$ values in the HP+ model. The PMF of the purely hydrophobic sequence ($X_{\mathrm{P}} = 0$; black dashed line) is shown as a reference in all subplots.

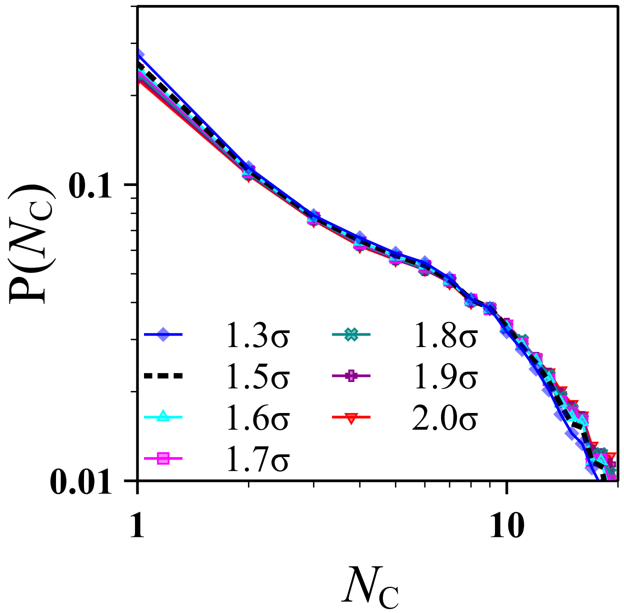


Figure S12: Probability of finding a chain in a cluster of size $N_c$ for $X_P = 0.75$ sequence in the HP+ model. Cluster sizes were determined based on different cutoff distances as denoted in the legend, with $\sigma = 0.5\,\text{nm}$ being the monomer diameter. Note that the cutoff distance $1.5\sigma$, shown as a dashed black line, was used to perform clustering analysis for different $X_P$ values of the primary sequence set in Fig. 7.

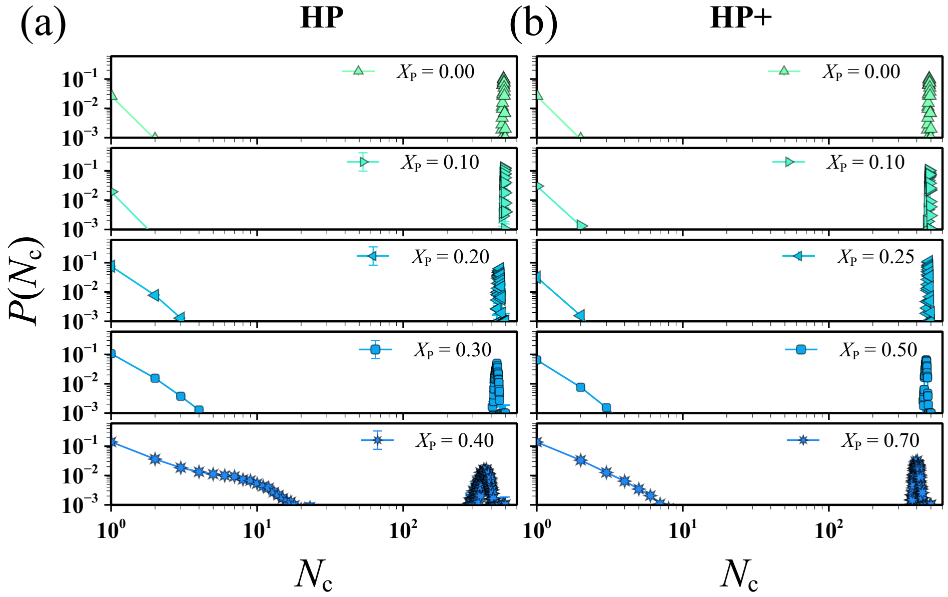


Figure S13: Probability of finding a chain in a cluster of size $N_c$ for select $X_P$ values that phase separated in the HP and HP+ models.

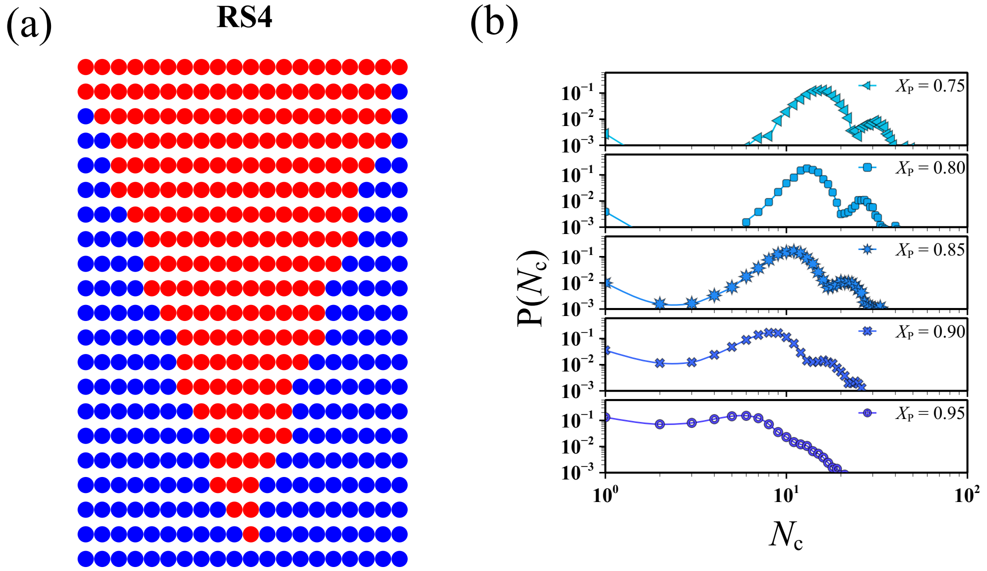


Figure S14: (a) Highly patterned sequence set (RS4) used to elucidate the effect of patterning on the formation of finite-sized aggregates. (b) Probability of finding a chain in a cluster of size $N_c$ for select $X_P$ values from RS4 that did not phase separate in the HP+ model.
